# Direct imaging of anthrax intoxication in animals reveals shared and individual functions of CMG-2 and TEM-8 in cellular toxin entry

**DOI:** 10.1101/2021.01.22.427304

**Authors:** Carly Merritt, Elizabeth M. Chun, Rasem J. Fattah, Mahtab Moayeri, Dennis Paliga, Sebastian Neumann, Rolf Heumann, Stephen H. Leppla, Thomas H. Bugge

## Abstract

The virulence of *Bacillus anthracis* is linked to the secretion of anthrax lethal toxin and anthrax edema toxin. These binary toxins consist of a common cell-binding moiety, protective antigen (PA), and the enzymatic moieties, lethal factor (LF) and edema factor (EF). PA binds either of two specific cell surface receptors, capillary morphogenesis protein-2 (CMG-2) or tumor endothelial marker-8 (TEM-8), which triggers the binding, endocytosis, and cytoplasmic translocation of LF and EF. The cellular distribution of functional TEM-8 and CMG-2 receptors during anthrax toxin intoxication in animals is not fully elucidated. Herein, we describe a novel assay to image anthrax toxin intoxication in live animals, and we use the assay to visualize TEM-8- and CMG-2-dependent intoxication. Specifically, we generated a chimeric protein consisting of the N-terminal domain of LF fused to a nuclear localization signal-tagged Cre recombinase (LFn-NLS-Cre). When PA and LFn-NLS-Cre were co-administered to transgenic mice that ubiquitously express a red fluorescent protein in the absence of Cre activity and a green fluorescent protein in the presence of Cre activity, anthrax toxin intoxication could be visualized at single-cell resolution by confocal microscopy. By using this assay, we show that CMG-2 is critical for intoxication in the liver and heart, whereas TEM-8 is required for full intoxication in the kidney and spleen. Other tissues examined were largely unaffected by single deficiences in either receptor, suggesting extensive overlap in TEM-8 and CMG-2 expression. The novel assay will be useful for basic and clinical/translational studies of *Bacillus anthracis* infection and for identifying on- and off-targets for reengineered toxin variants in the clinical development of cancer treatments.

**Background:** Assays for imaging of anthrax toxin intoxication in animals are not available.

**Results:** Anthrax toxin-Cre fusions combined with fluorescent Cre reporter mice enabled imaging of anthrax toxin intoxication in animals.

**Conclusion:** Shared and distinct functions of toxin receptors in cellular entry were uncovered. Significance. A simple and versatile assay for anthrax toxin intoxication is described.

---

Anthrax is contracted through inhalation, ingestion, or cutaneous inoculation of endospores of the Gram-positive bacterium *Bacillus anthraci*s. Spores germinate following their introduction to the body and disseminate to cause a systemic infection, which, if left untreated, is associated with high mortality rates. Upon the death of the host, *Bacillus anthracis* forms spores that are resistant to chemical insults, heat exposure, and dehydration and remain infectious for long periods (1,2).

The virulence of *Bacillus anthracis* results from the release of three proteins into the circulation: protective antigen (PA), lethal factor (LF), and edema factor (EF). These three proteins are individually nontoxic, and PA combines with either LF to form anthrax lethal toxin or with EF to form anthrax edema toxin. The systemic administration of anthrax toxin to animals closely mimics experimental infection with *Bacillus anthraci*s, and vaccination against the toxin components is protective, indicating that anthrax is largely a toxin-mediated disease (1,2). Anthrax toxins exert their cytotoxic actions in a three-step activation process that involves: a) the binding of PA to the surface of target cells, b) the translocation of LF and EF to the cytoplasmic compartment of the target cells, and c) the enzymatic action of LF and EF on cytoplasmic substrates (1,2). Anthrax toxin intoxication is initiated by the binding of PA to either of two receptors, capillary morphogenesis protein-2 (CMG-2) or tumor endothelial marker-8 (TEM-8) (3,4). Subsequently, PA is cleaved at the sequence, ^164^RKKR^167^, by cell surface-localized furin or furin-like pro-protein convertases (5). This endoproteolytic cleavage is absolutely required for toxin activation and triggers all subsequent steps of the activation process. The C-terminal 63 kDa fragment of PA (PA63) remains bound to the cell surface after endoproteolytic cleavage and undergoes a conformational change that leads to the formation of a PA63 heptamer or octamer that subsequently binds up to four molecules of LF or EF with high affinity (6–8). The complex of PA63 and LF or EF is then endocytosed, and PA63 undergoes pH-induced conformational changes in the endosomal/lysosomal compartment to form a channel that facilitates the unfolding and translocation of LF and EF to the cytoplasm. EF is an adenylate cyclase proposed to lead to the formation of supraphysiological intracellular levels of cyclic AMP (9). LF is a zinc-dependent metalloproteinase that can cleave and inactivate several mitogen-activated protein kinase kinases (10). Although essential for intoxication, the cellular distribution of CMG-2 and TEM-8 and the function of each receptor in the intoxication in specific organs remain to be fully elucidated. Notably, assays for direct visualization of anthrax toxin intoxication *in vivo* are not available, and the tissue and cellular targets for anthrax toxin during *in vivo* infection have been inferred only indirectly from analysis of tissues from intoxicated animals or from biochemical and genetic analysis of anthrax toxin targets (11,12).

LF is stable in circulation when administered alone and only becomes cell surface-associated after the binding of PA to CMG-2 or TEM-8 and its subsequent proteolytic cleavage to PA63. It has long been noted that LF residues 1-254 suffice to achieve translocation of a variety of “passenger” polypeptides and other molecules into the cytoplasm of the cells in a PA63-dependent manner (13,14). These include other bacterial toxins and bacterial proteins (13,15–19), fluorescent proteins (20), viral proteins (21–24), eukaryotic proteins (25–28), and radioisotopes (29–32). Thus, the fusion or conjugation of LF to imageable moieties could provide ideal agents for studying the cellular intoxication by anthrax toxin *in vivo*. A significant caveat to this approach, however, is the low number of LF molecules successfully translocated to the cytoplasm through the PA pore, which makes most imaging modalities poorly suited to study anthrax toxin intoxication in whole-animal systems (18,27). A second challenge to whole-animal imaging approaches is that most imaging modalities, such as radionuclides, enzymes such as horseradish peroxidase and βgalactosidase, and fluorescent proteins, likely would not discriminate between productive intoxication (i.e. PA-dependent entry into the cytoplasm) and non-productive interactions of the labeled toxin with cells, such as cell surface retention, fluid phase pinocytosis, and endosomal/lysosomal accumulation of intact or partially degraded toxin conjugates.

Spleen extracts from reporter mice carrying a Cre-activated βgalactosidase transgene have been shown to express increased β-galactosidase activity when infected with *Salmonella enterica* serovar Typhimurium carrying the type III secreted protein, SopE, fused to bacteriophage P1 Cre recombinase (33). Although single-cell resolution was not achieved, presumably due to a low number of cells being infected (33), the study provided evidence that bacterial protein-Cre fusion proteins may display sufficient enzymatic activity in animals to induce LoxP-dependent recombination.

In this study, we used a combined biochemical and genetic approach to image anthrax toxin intoxication in animals. Specifically, we generated a tripartite fusion protein that consists of the N-terminal domain of LF fused to a nuclear localization signal-tagged bacteriophage P1 Cre recombinase (LFn-NLS-Cre). When PA and LFn-NLS-Cre were co-administered to transgenic mice that ubiquitously express a red fluorescent protein (tdTomato) in the absence of Cre activity and a green fluorescent protein (eGFP) in the presence of Cre activity (hereafter *mTmG* mice), anthrax toxin intoxication could readily be visualized at single-cell resolution by using confocal microscopy of unfixed and unprocessed organs. By superimposing individual genetic deficiencies of either TEM-8 or CMG-2 in *mTmG* mice, we were able to directly establish the importance of each receptor in anthrax toxin intoxication in individual tissues.

The assay presented here should be useful for basic and clinical/translational studies of *Bacillus anthracis* infection, and it may be easily adapted to image intoxication in animals by related bacterial type III toxins. Furthermore, the assay will be valuable for identifying on- and off-targets for reengineered anthrax toxins currently in clinical and pre-clinical development for the treatment of solid tumors (34–50). When used in conjunction with modified PA variants that are activated by specific cell surface proteases, the assay may also be suitable for *in vivo* imaging of specific cell surface proteolytic activity in a variety of physiological and pathological processes.

## EXPERIMENTAL PROCEDURES

### Recombinant proteins

Plasmids for expressing proteins having LFn (LF aa 1-254) fused to Cre recombinase were constructed using the Champion pET SUMO vector (Invitrogen, Carlsbad, CA), which expresses proteins fused at the C-terminus of a His6-SUMO tag. DNA-encoding residues 1-254 of anthrax lethal factor originated from *Bacillus anthracis*, and those of Cre recombinase from bacteriophage P1. The sequences of the four LFn-Cre proteins are shown in Supplemental Figure 1. They differ only in the linker sequences. Two of the constructs contain ubiquitin between LFn and Cre, and three contain a nuclear localization signal (NLS) preceding Cre. The proteins were expressed as per the Champion pET SUMO manufacturer’s instructions with minor modifications. Cell lysates were adsorbed to Ni-NTA resins, and the His-tagged proteins were eluted with imidazole solutions. The His6-SUMO-LFn-Cre proteins were cleaved with SUMO protease, which was made inhouse using Addgene plasmid #64697, deposited to Addgene (Watertown, MA) by Dr. Hideo Iwai (51). The His6-SUMO polypeptide and the His6-tagged SUMO protease were subtractively removed by passage through Ni-NTA resin. The LFn-Cre proteins were further purified by chromatography on hydroxyapatite to achieve purities of >95%. The LFn-NLS-Cre protein selected for the animal imaging studies was obtained in yields of >20 mg/L of culture.

**Figure 1.**
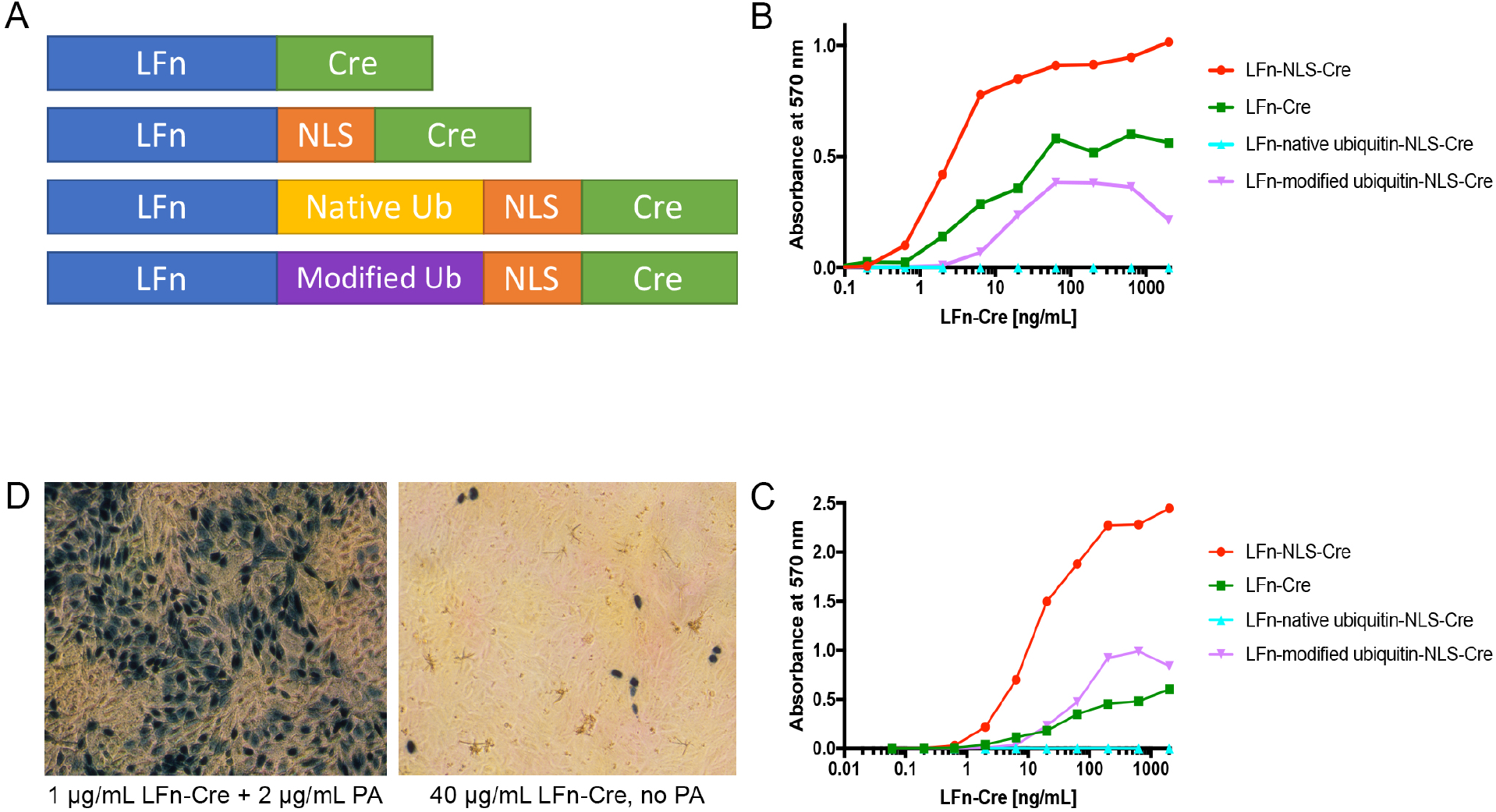
Generation of LFn-Cre fusion proteins capable of PA-dependent cellular entry. A. Schematic structure of LFn-Cre, LFn-NLS-Cre, LFn-native ubiquitin-NLS-Cre, and LFn-modified ubiquitin-NLS-Cre fusion proteins. B and C. CV1-5B cells, harboring a Cre-activated β-galactosidase gene, were plated at low density (B) or high density (C) and incubated with increasing concentrations of LFn-NLS-Cre, LFn-Cre, LFn-native ubiquitin-NLS-Cre, or LFn-modified ubiquitin-NLS-Cre fusion proteins in the presence of 1 βg/mL PA. β-galactosidase activity was measured after 40 h by using the chromogenic substrate, chlorophenolred-β-D-galactopyranoside, and was plotted against the LFn-Cre fusion protein concentration. D. CV1-5B cells were incubated with 1 βg/mL LFn-NLS-Cre and 2 βg/mL PA (left panel) or with 40 βg/mL LFn-NLS-Cre alone (right panel). The cells were stained *in situ* for β-galactosidase activity (blue) after 40 h.

### Cell culture assays

Efficacy of LFn-Cre protein translocation into cells was tested in CV1-5B cells (52,53). Cells were plated in 96-well plates in DMEM high glucose medium with 10% fetal bovine serum, cultured at 37°C at 10% CO_2_, and used when they were at low and high confluency. PA was added at 1 μg/mL in all wells, and LFn-Cre proteins were diluted serially at 3.14-fold. After 40 h, cells were washed in phosphate buffered saline (PBS) containing 2 mM MgCl_2_ and fixed in PBS, 5 mM EGTA, 2 mM MgCl_2_, and 0.2% glutaraldehyde for 30 min. After again washing in PBS with 2 mM MgCl_2_, the cells were stained for β-galactosidase activity with PBS, 2 mM MgCl_2_, 0.1% Triton X-100, 0.1% NaN_3_, and 1 mg/mL chlorophenolred-β-D-galactopyranoside. Absorbance was measured at 570 nm after 20 min to quantify conversion of substrate by β-galactosidase (54).

### Animal work

All experiments were performed in an Association for Assessment and Accreditation of Laboratory Animal Care International-accredited vivarium following Standard Operating Procedures and were approved by the NIAID Institutional Animal Care and Use Committee. B6.129(Cg)-*Gt(ROSA)26Sortm4(ACTB-tdTomato,-EGFP)Luo*/J^+/0^ (*mTmG*^+/0^ mice) (55) were generated by crossing B6.129(Cg)-*Gt(ROSA)26Sortm4(ACTB-tdTomato,-EGFP)Luo*/J^+/+^ (Jackson Laboratory, Bar Harbor, ME) males with C57BL/6J females (Jackson Laboratory, Bar Harbor, ME). Wildtype C57BL/6J mice were used as controls. Both male and female mice were used for experiments at age 4-8 weeks. The generation of CMG-2-(*Cmg2*^-/-^) and TEM-8-(*Tem8*^-/-^) deficient mice has been described previously (56). *Cmg2*^-/-^; *mTmG*^+/0^ and *Tem8*^-/-^;*mTmG*^+/0^ mice were generated by interbreeding. The following primer pairs were used for genotyping: *Cmg2* wildtype allele, 5’-GACTCTTAGGAAGGGTTCCTACTGG-3’ and 5’-TGTAAGTCATATGGGTAGTGACCTAT-3’. *Cmg2* mutant allele, 5’-CCAATTTGGAGCTCAGGTTGGTGGA-3’ and 5’-TGTAAGTCATATGGGTAGTGACCTAT-3’. *Tem8* wildtype allele, 5’-ATGTCCCTTTGCCTCTTGTGGC-3’ and 5’-TTCCACCTCACTGACCACCC-3’. *Tem8* mutant allele, 5’-AGGCACTGACAAACCCTCTCAGGA-3’ and 5’-CCAGCCCATGCTGACAGCTCACAGA-3’.

### PCR analysis of Cre-mediated recombination in mouse organs

25 µg of LFn-NLS-Cre and 25 µg PA in PBS were delivered via tail vein injection into *mTmG*^+/0^ mice. 24 h post injection, the mice were euthanized by CO_2_ inhalation. Organs and bone marrow were digested overnight in lysis buffer (100 mM Tris HCl, pH 8, 5 mM EDTA, 200 mM NaCl, 0.2% SDS, and 100 mg/mL Proteinase K) for 24 h at 55^°^C. DNA was precipitated using isopropanol. PCR was performed with the following primers: GFP RV: 5’-CGTCGCCGTCCAGCTCGACCAG-3’, Tomato RV: 5’-GCCCATGAACTCTTTGATGACCTCCTCTCCC-3’, and LoxP FW: 5’-CCGCGGGCTCGACACTGAACC-3’ using JumpStart REDTaq ReadyMix (Sigma, St Louis, MO). PCR products were separated on 1.5% agarose gels, stained with ethidium bromide, and visualized by UV illumination. Cre recombination resulted in a PCR product (LoxP FW + GFP RV) of 258 bp, while the non-recombinant PCR product (LoxP FW + Tomato RV) was 157 bp.

### Imaging anthrax toxin intoxication in mice

LFn-NLS-Cre and PA proteins alone or in combination in PBS were delivered intraperitonially or via tail vein injection. The mice were tail vein-injected with 100 μL of 6 mg/mL Hoechst dye (Thermo Fisher Scientific, Waltham, MA) 4-6 h prior to termination of an experiment to visualize nuclei (57). Mice were euthanized by CO_2_ inhalation and perfused with ice-cold PBS using cardiac puncture. Organs were immediately removed and cut into ∼1-2 mm thick slices using a scalpel. The organ slices were placed on a MatTek glass bottom microwell dish (MatTek Corporation, Ashland, MA) and imaged using a 20x 0.75 NA Air or 60x 1.27 NA Water objective (Nikon, Tokyo, Japan) on an A1R + MP confocal microscope system (Nikon, Tokyo, Japan). Large images were composed of stitched images with a 10% overlap using NIS-Elements software (Nikon, Tokyo, Japan).

## RESULTS

### Generation of LFn-Cre recombinase fusion proteins capable of PA-dependent cytoplasmic translocation

We have previously shown that PA-dependent translocation of a LF-β-lactamase fusion protein can be imaged in cultured cells by using a cell-penetrating β-lactamase quenched fluorescence resonance energy transfer substrate (18). The adaptation of this assay for imaging anthrax toxin intoxication in whole animals, while hypothetically feasible, is prohibited by the high cost of the β-lactamase substrate and, likely, by logistic problems associated with systemic delivery of the substrate to animals. We therefore explored the possibility of combining biochemical and genetic approaches to imaging anthrax toxin intoxication in whole animals. Specifically, we generated a series of proteins that consist of the PA-binding domain of LF (LFn) either fused directly to the bacteriophage P1 Cre recombinase (LFn-Cre) or linked via a nuclear localization signal (LFn-NLS-Cre), a native ubiquitin protein (LFn-native ubiquitin-NLS-Cre), or a modified ubiquitin protein followed by a nuclear localization signal (LFn-modified ubiquitin-NLS-Cre) (Figure 1A). The inclusion of ubiquitin as a linker was based on the expectation that ubiquitin fusions of this type, once delivered to the cytosol, would be cleaved by deubiquitinating enzymes, releasing the Cre domain as a smaller entity that may have greater access to the nucleus, the location of its target DNA substrate (58). Furthermore, release from LFn could decrease the probability that ubiquitination of the LFn region would deliver the entire fusion protein to the proteasome for degradation. Similarly, the replacement of the seven Lys residues of ubiquitin with Arg in LFn-modified ubiquitin NLS-Cre could limit ubiquitination and increase stability.

Preliminary testing of the four purified LFn-Cre proteins was done in CV1-5B reporter cells, which contain a Cre-activated β-galactosidase gene, by treating the cells with increasing concentrations of fusion protein in the presence of a fixed concentration of PA (Figure 1B and C). This experiment identified LFn-NLS-Cre as the most effective of the four fusion proteins. This difference was most prominent in cells plated at high density (compare Figure 1B and C). The need for an NLS, the only feature present in LFn-NLS-Cre compared to LFn-Cre (Figure 1A), suggests that the NLS reported to exist in Cre (59) is not by itself sufficient to maximize nuclear delivery in this context. The alternative explanation, that the added NLS in LFn-NLS-Cre provides a target by which cytosolic proteases release the Cre domain, cannot be discounted without further study. The total inactivity of LFn-native ubiquitin-NLS-Cre compared to that of LFn-modified ubiquitin-NLS-Cre indicated that native ubiquitin is unable to translocate through the PA pore. The entry of LFn-NLS-Cre into cells was highly PA-dependent, with very few CV1-5B cells displaying β-galactosidase activity in the absence of PA, even when LFn-NLS-Cre was administered at a 40-fold higher concentration (Figure 1D).

### Imaging anthrax toxin intoxication in mice

The above studies showed that LFn-Cre fusion proteins could translocate to the cytoplasm in a PA-dependent manner, that the Cre moiety (alone or as an intact fusion protein) thereafter was imported to the nucleus, and that it retained its recombinase activity after nuclear translocation. This indicated that LFn-Cre fusion proteins, when used in conjunction with appropriate transgenic mouse Cre reporter strains, could be suitable for imaging anthrax toxin intoxication *in vivo*. The B6.129(Cg)-*Gt(ROSA)26Sortm4(ACTB-tdTomato,-EGFP)Luo*/J transgenic mouse (hereafter *mTmG*) is a widely used reporter strain for Cre activity (55). The strain constitutively expresses a floxed plasma membrane-tagged tdTomato (mTomato) fluorescent protein gene under the control of a cytomegalovirus-β-actin enhancer-promoter. This gene is excised by Cre, which simultaneously allows for transcription of a plasma membrane-tagged enhanced green fluorescent protein (eGFP)-expressing gene by placing it proximal to the cytomegalovirus-β-actin enhancer-promoter. Thus, following cytoplasmic translocation of LFn-NLS-Cre through the PA pore and subsequent nuclear import, intoxicated cells should display green fluorescent plasma membranes, while plasma membranes of non-intoxicated cells should display only red fluorescence.

To test if PA and LFn-NLS-Cre could mediate LoxP-dependent recombination in a whole-animal context, we first designed PCR primer sets that would selectively amplify, respectively, the non-recombined and the recombined *mTmG* transgene (Figure 2A). Interestingly, a PCR product derived from the recombined transgene was readily detected in the heart, lungs, liver, kidney, spleen, lymph nodes, thymus, uterus, esophagus, trachea, tongue, and bone marrow of *mTmG*^+/0^ mice injected with LFn-NLS-Cre and PA but not in these organs from non-injected *mTmG*^+/0^ mice (Figure 2B). This PCR product was not observed in the intestine, skin, and brain of mice injected with LFn-NLS-Cre and PA (Figure 2B).

**Figure 2.**
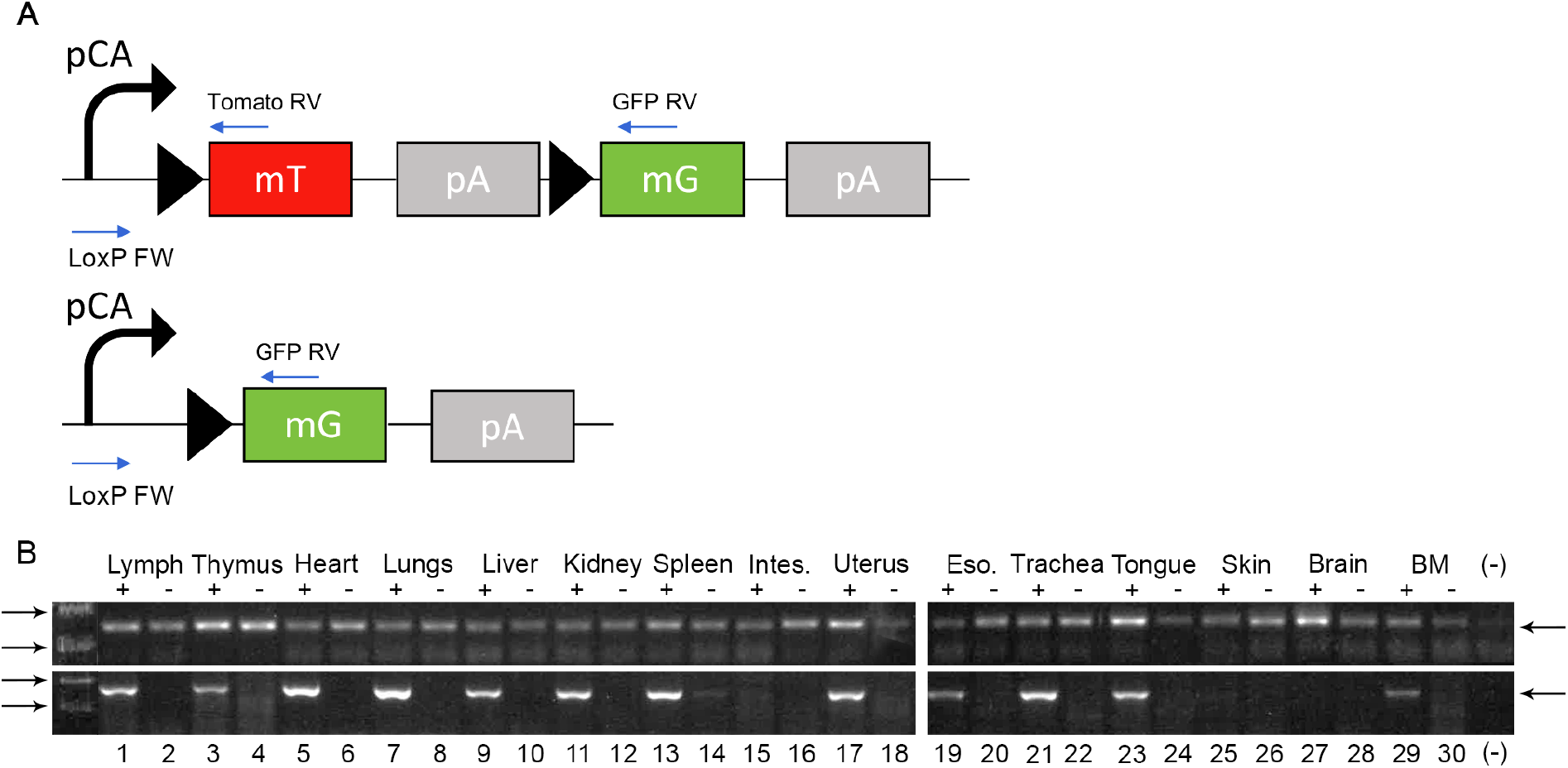
Analysis of LFn-NLS-Cre activity in toxin-injected *mTmG*^+/0^ mice by PCR. A. Schematic depiction of *mTmG* transgene and location of primers used for detection of non-recombined (LoxP FW and Tomato RV) and recombined (LoxP FW and GFP RV) alleles. The positions of LoxP sites are indicated. B. Ethidium bromide-stained agarose gels of PCR products generated from DNA isolated from lymph nodes (lanes 1 and 2), thymus (lanes 3 and 4), heart (lanes 5 and 6), lungs (lanes 7 and 8), liver (lanes 9 and 10), kidney (lanes 11 and 12), spleen (lanes 13 and 14), intestine (Intes, lanes 15 and 16), uterus (lanes 17 and 18), esophagus (Eso, lanes 19 and 20), trachea (lanes 21 and 22), tongue (lanes 23 and 24), skin (lanes 25 and 26), brain (lanes 27 and 28), and bone marrow (BM, lanes 29 and 30) of an *mTmG*^+/0^ mouse injected with 25 μg LFn-NLS-Cre and 25 µg PA (lanes 1, 3, 5, 7, 9, 11, 13, 15, 17, 19, 21, 23, 25, 27, and 29) or a non-injected *mTmG*^+/0^ littermate (lanes 2, 4, 6, 8, 10, 12, 14, 16, 18, 20, 22, 24, 26, 28, and 30). Positions of PCR products expected to be generated from non-recombined (157 bp) and recombined (258 bp) *mTmG* alleles are shown with arrows at right. Positions of molecular weight markers (100 and 200 bp for top panel and 200 and 300 bp for bottom panel) are shown at left. (-), no DNA.

We next determined the ability to detect fluorescence in *mTmG*^+/0^ mice by confocal microscopy of unfixed whole organ slices, which could serve as a convenient readout for Cre activity. Wildtype mice were analyzed in parallel as a control for autofluorescence. To obtain semi-quantitative estimates of fluorescence intensities of the examined organs, in this and the following experiments, we generated composite images of the entire organ slice from confocal images acquired at low magnification. As expected, red fluorescence of variable intensity was observed in multiple organs of *mTmG*^+/0^ mice but not in the corresponding organs from wildtype mice imaged using the identical conditions (Supplemental Figure 2). This indicated that the *mTmG* mouse strain, in principle, should be suitable for the detection of anthrax toxin intoxication in cells in a variety of organs and tissues by this simple procedure.

Having established that fluorescence in *mTmG*^+/0^ mice can readily be detected by confocal microscopy of unfixed whole organ slices, we next intravenously injected *mTmG*^+/0^ and wildtype mice with LFn-NLS-Cre alone, PA alone, or with PA in combination with LFn-NLS-Cre (Supplemental Figure 3). Mice were killed 24 h later, and slices of excised organs were immediately examined by confocal microscopy. Organs from *mTmG*^+/0^ mice injected with LFn-NLS-Cre alone displayed only red fluorescence (Supplemental Figure 3 A, A’, D, D’, G, G’, J, J’, M, M’), importantly showing that the LFn-NLS-Cre protein by itself does not gain access to cells. As expected, organs from mice injected with PA alone also displayed only red fluorescence (Supplemental Figure 3 B, B’, E, E’, H, H’, K, K’, N, N’). Interestingly, however, when analyzing these organs from *mTmG*^+/0^ mice injected with LFn-NLS-Cre in combination with PA, both red (Supplemental Figure 3 C, F, I, L, O) and green fluorescence were readily detected in the heart, kidney, liver, lungs, and spleen (Supplemental Figure 3 C’, F’, I’, L’, O’), indicating that PA-dependent translocation of LFn-NLS-Cre to the cytoplasm of cells in these organs takes place and that the subsequent nuclear import of the fusion protein leads to the excision of the floxed mTomato gene. No green fluorescence was observed in the same organs of wildtype mice injected with LFn-NLS-Cre in combination with PA, further demonstrating the origin of the green fluorescent signal from *de novo* expression of eGFP (Supplemental Figure 4, compare A with B, C with D, E with F, G with H, I with J). Compatible with the PCR analysis, green fluorescence was weak or absent in intestine (Supplemental Figure 5 A, B), skin (Supplemental Figure 5 C, D), and brain (Supplemental Figure 5 E, F) slices, although cells in these organs of *mTmG*^+/0^ mice were previously shown to undergo recombination *in vivo* and express eGFP in the presence of a Cre-expressing transgene (55).

### Effect of route of administration, dose, and time of analysis on imaging of intoxication

Parameters affecting the intoxication process and the optimal use of mTomato/eGFP fluorescence as a readout for anthrax toxin intoxication would be expected to include the mode and dose of toxin delivery, tissue accessibility, receptor expression, efficiency of cytoplasmic translocation and nuclear transport of the LFn-NLS-Cre fusion protein, rate of Cre-mediated recombination, decay rate of the mTomato mRNA and protein, the rate of transcription, translation and attainment of steady state levels of the eGFP protein, ploidy of the target cells (diploid versus polyploid, carrying one or more than one transgene), and time of imaging after toxin administration. The next set of experiments were aimed at empirically addressing some of these issues by varying the mode and dose of toxin delivery, number of injections, and time from injection to imaging. We first compared low-magnification confocal images of green fluorescence obtained from whole organ slices of the heart, kidney, liver, lungs, and spleen from *mTmG*^+/0^ mice injected either intravenously or intraperitoneally with PA in combination with LFn-NLS-Cre (Supplemental Figure 6). This analysis showed that toxin administered intraperitoneally intoxicated the organs less efficiently, as evidenced by less green fluorescence, than when administered intravenously (Supplemental Figure 6, compare A-C with D, E-G with H, I-K with L, M-O with P, Q-S with T).

We next studied the effect of multiple injections of PA and LFn-NLS-Cre on the intoxication in the five organs analyzed above (Supplemental Figure 7). Specifically, we injected *mTmG*^+/0^ mice with PA and LFn-NLS-Cre at 0 h, 24 h, 48 h, and 72 h and procured low-magnification red and green confocal fluorescence images of organ slices at 24 h, 48 h, 72 h, and 120 h. For each time point analyzed, a separate control group of mice injected with PA and LFn-NLS-Cre once at 0 h was included to control for Cre-mediated recombination occurring during the assay period and/or the eGFP protein not having attained steady state levels at 24 h. This control experiment, indeed, showed that the eGFP signal in organs of mice having received a single injection of PA and LFn-NLS-Cre increased qualitatively beyond 24 h, with the possible exception of the liver (Supplemental Figure 7, compare A’ with B’, C’, and D’, E’ with F’, G’, and H’, I’ with J’, K’, and L’, M’ with N’, O’, and P’, Q’ with R’, S’, and T’). However, for the kidney (Supplemental Figure 7, compare F’ with F’’’, G’ with G’’’, H’ with H’’’), liver (Supplemental Figure 7, compare J’ with J’’’, K’ with K’’’, L’ with L’’’), and spleen (Supplemental Figure 7, compare R’ with R’’’, S’ with S’’’, T’ with T’’’), an increased eGFP signal was evident in mice receiving multiple injections of PA and LFn-NLS-Cre, as compared to single-injected control mice. Interestingly, the mTomato signal in the liver was greatly diminished already at 48 h in mice receiving either single or multiple injections of PA and LFn-NLS-Cre (Supplemental Figure 7, compare I with J and J’’). This indicates that intoxication in the majority of cells in this organ was achieved.

To examine the effect of the dose of toxin administered on intoxication, we injected *mTmG*^+/0^ mice with 5, 15, and 25 μg PA and LFn-NLS-Cre and imaged their organs 24 h later. At the 5 μg dose, only the lungs displayed a faint eGFP signal (Supplemental Figure 8 J’). At the 15 μg dose, eGFP signals were seen in the liver (Supplemental Figure 8 H’) and lungs (Supplemental Figure 8 K’). At the 25 μg toxin dose, an eGFP signal also became evident in the heart (Supplemental Figure 8 C’), kidney (Supplemental Figure 8 F’), and spleen (Supplemental Figure 8 O’).

To determine when an eGFP signal is first detectable after the administration fo PA and LFn-NLS-Cre, we injected *mTmG*^+/0^ mice with 75 μg of each protein and examined the heart, kidney, liver, lungs, and spleen by confocal microscopy at 6, 8, 10, and 12 h (Supplemental Figure 9). Whereas no signal was observed at any of these time points in the heart (Supplemental Figure 9 A’, B’, C’, D’), kidney (Supplemental Figure 9 E’, F’, G’, H’), lungs (Supplemental Figure 9 M’, N’, O’, P’), and spleen (Supplemental Figure 9 Q’, R’,S’, T’), faint eGFP signals were observed in the liver at 12 h (Supplemental Figure 9 L’).

### Single-cell resolution imaging of anthrax toxin intoxication

Using the knowledge gained from the above experiments, we next tested the ability of the assay to image intoxication in individual cells in unprocessed organs. For this purpose, mice received three intravenous injections of 25 µg LFn-NLS-Cre and 25 µg PA at 0 h, 24 h, and 48 h. 72 h after the first injection, confocal images of red, green, and blue (nuclei) fluorescence of slices of the heart, kidney, liver, lungs, and spleen were acquired at high magnification (Figure 3). This analysis showed that in tissues of these five organs, non-intoxicated and intoxicated individual cells were readily distinguishable by their red and their green or yellow membrane-confined fluorescence, respectively.

**Figure 3.**
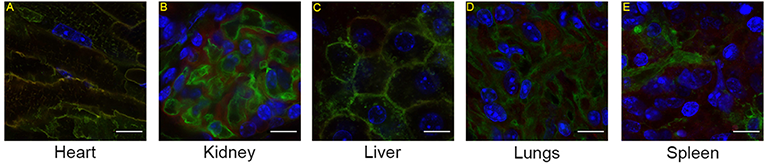
Imaging of anthrax toxin intoxication of individual cells. High-magnification confocal images of slices of the heart (A), kidney (B), liver (C), lungs (D), and spleen (E) from *mTmG*^+/0^ mice after intravenous administration of 25 µg LFn-NLS-Cre and 25 µg PA at 0 h, 24 h, and 48 h with analysis at 72 h. Intoxication in individual cells of cardiac muscle, kidney glomerulus, liver parenchyma, lung alveoli, and spleen is revealed by membrane-associated green or yellow staining. Nuclei (blue) were visualized by systemic Hoechst administration to the mice prior to analysis. The images were collected using a 60x objective. Size bars: 10 µm.

### Effect of genetic elimination of CMG-2 and TEM-8 on anthrax toxin intoxication

We next interbred previously generated CMG-2-deficient (*Cmg2*^-/-^) and TEM-8-deficient (*Tem8*^-/-^) mice to *mTmG*^+/0^ mice to generate, respectively, *Cmg2*^-/-^;*mTmG*^+/0^ and *Tem8*^-/-^;*mTmG*^+/0^ mice. These mice, together with *mTmG*^+/0^ mice, were injected at 0 h, 24 h, 48 h, and 72 h with 25 µg LFn-NLS-Cre and 25 µg PA, and red and green confocal fluorescence images of organ slices from the heart, kidney, liver, lungs, spleen, intestine, and skin were acquired at 120 h. This injection scheme, as expected from the preceeding experiments, resulted in green fluorescence of cells from the heart, liver and lungs, kidney, and spleen of *mTmG*^+/0^ mice (Figure 4A-A’’’, 4C-C’’’, 4E-E’’’, 4G-G’’’). In sharp contrast, green fluorescent cells were essentially absent in the heart and liver of *Cmg2*^-/-^;*mTmG*^+^ mice, showing that CMG-2 mediates intoxication in these organs uncompensated by TEM-8 (Figure 4 B-B’’’, F-F’’’). The pattern of fluorescence in the kidney, lungs, and spleen was not appreciably different between *mTmG*^+^ and *Cmg2*^-/-^;*mTmG*^+^ mice, revealing that CMG-2 is not critical for the intoxication in these organs in the presence of TEM-8 (Figure 4 D-D’’’, H-H’’’, J-J’’’).

**Figure 4.**
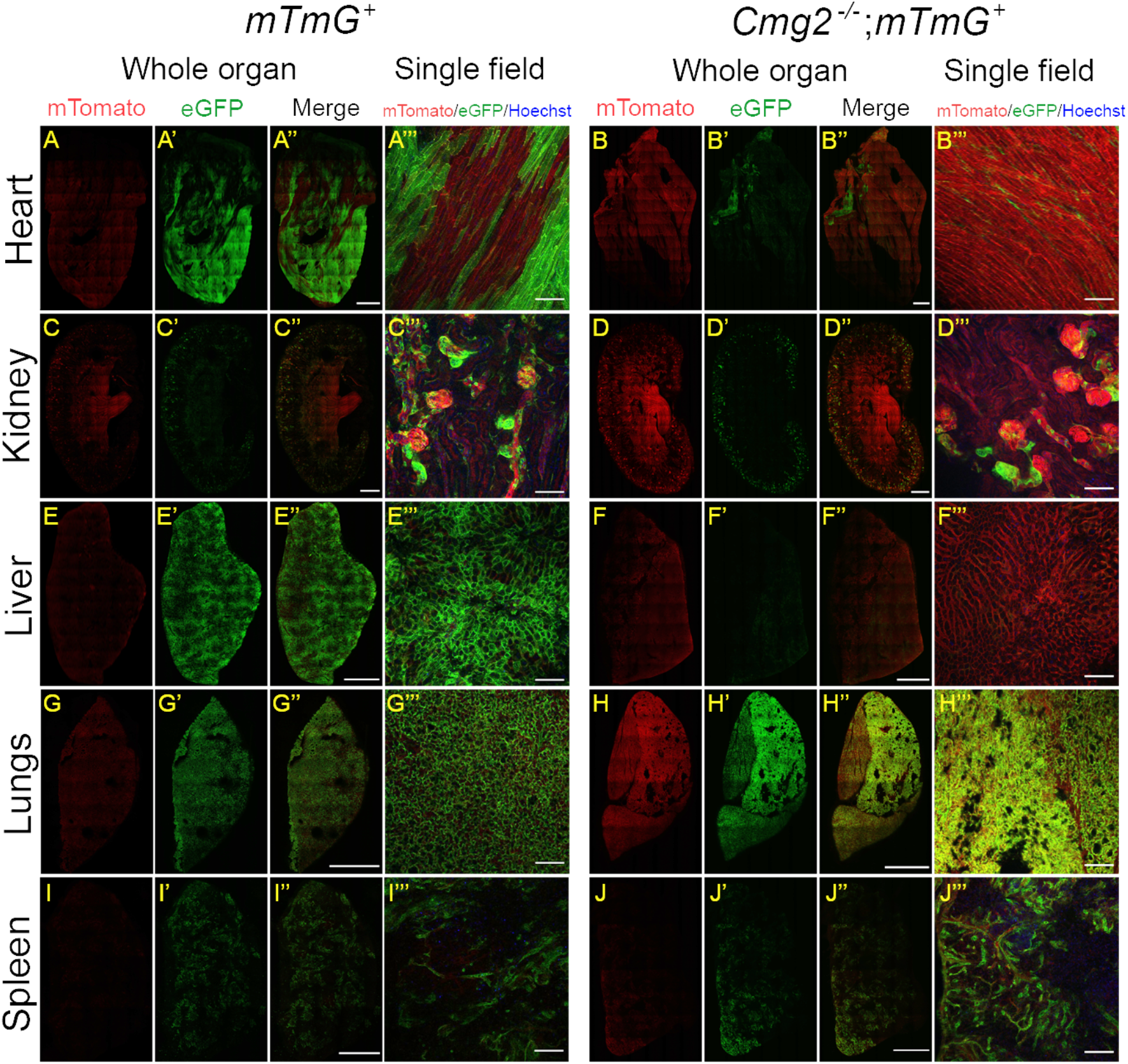
CMG-2 is critical for anthrax toxin intoxication in the heart and liver. Low-magnification confocal images of fresh slices of the heart (A-A’’’, B-B’’’), kidney (C-C’’’, D-D’’’), liver (E-E’’’, F-F’’’), lungs (G-G’’’, H-H’’’), and spleen (I-I’’’, J-J’’’) from CMG-2-sufficient, *mTmG-*sufficient (left panel) and CMG-2-deficient, *mTmG-*sufficient (right panel) mice after intravenous administrations of 25 µg LFn-NLS-Cre and PA at 0 h, 24 h, 48 h, 72 h, and 96 h with analysis at 120 h. The images were collected using a 20x objective. The whole organ images were assembled from individual images and were stitched together to cover the entire organ slice. Nuclei (blue) were visualized by systemic Hoechst administration to the mice prior to analysis. Size bars for whole organ: 1 mm. Size bars for single field: 100 µm. The data are representative of four separate experiments.

We next performed a similar comparison of toxin-treated *mTmG*^+^ and *Tem8*^-/-^;*mTmG*^+^ mice analyzed in parallel. This revealed that, unlike CMG-2, TEM-8 was not critical for the intoxication in the heart and liver (Figure 5B-B’’’, F-F’’’), revealing CMG-2 as the dominant receptor for the intoxication in these organs. Intoxication in the lungs was not substantially affected by the loss of TEM-8 (Figure 5H-H’’’), suggesting that intoxication in this organ is mediated by either of the two anthrax toxin receptors. In sharp contrast, the loss of TEM-8 markedly reduced the green fluorescent signal in the kidney (Figure 5 D-D’’’) and spleen (Figure 5J-J’), suggesting that TEM-8 plays a key role in the intoxication in these tissues.

**Figure 5.**
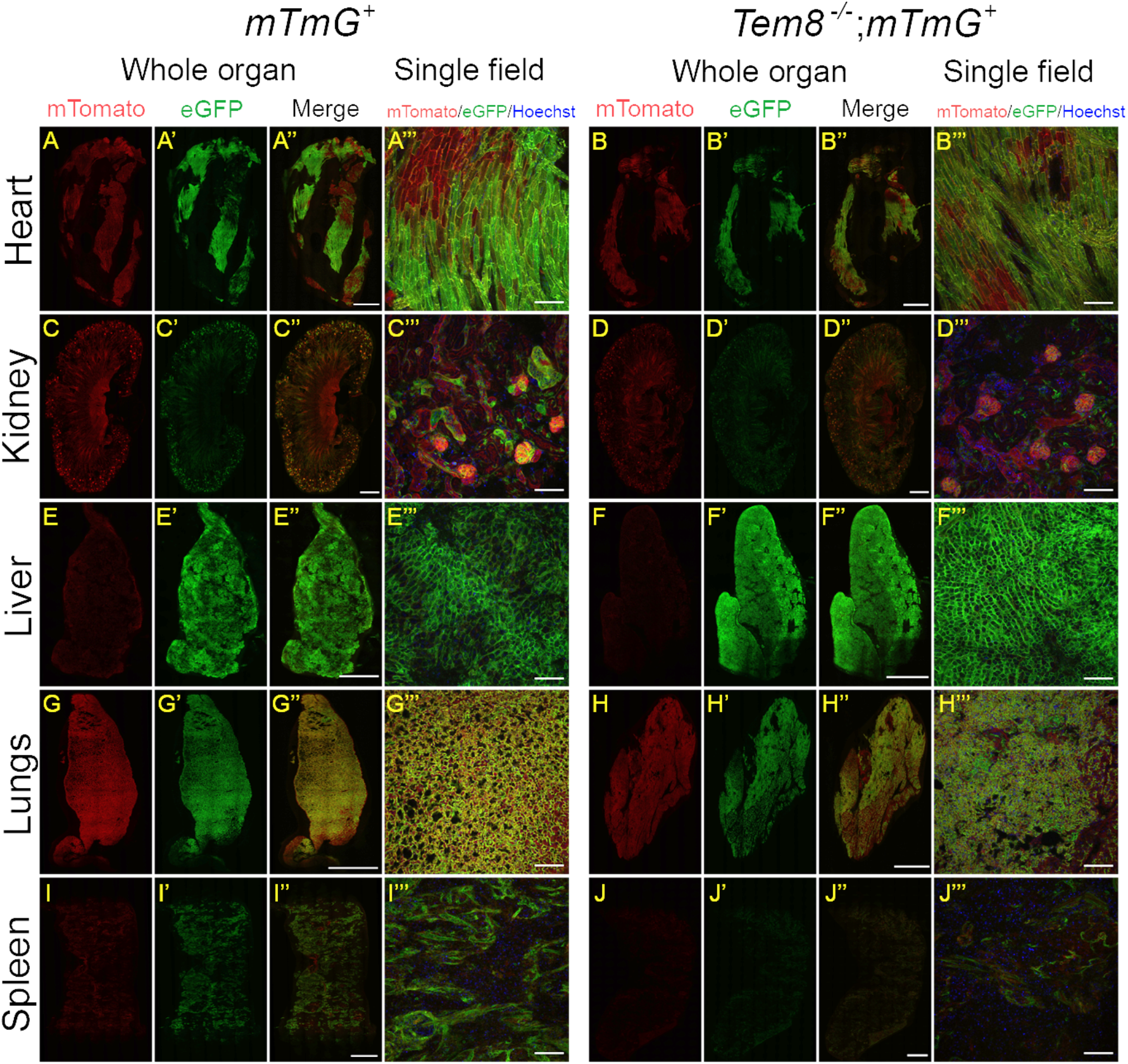
TEM-8 has a key role in anthrax toxin intoxication in the kidney and spleen. Low-magnification confocal images of fresh slices of the heart (A-A’’’, B-B’’’), kidney (C-C’’’, D-D’’’), liver (E-E’’’, F-F’’’), lungs (G-G’’’, H-H’’’), and spleen (I-I’’’, J-J’’’) from TEM-8-sufficient, *mTmG-*sufficient (left panel) and TEM-8-deficient, *mTmG-*sufficient (right panel) mice after intravenous administrations of 25 µg LFn-NLS-Cre and PA at 0 h, 24 h, 48 h, 72 h, and 96 h with analysis at 120 h. The images were collected using a 20x objective. The whole organ images were assembled from individual images and were stitched together to cover the entire organ slice. Nuclei (blue) were visualized by systemic Hoechst administration to the mice prior to analysis. Size bars for whole organ: 1 mm. Size bars for single field: 100 µm. The data are representative of four separate experiments.

## DISCUSSION

The ability to directly visualize the intoxication in cells by bacterial exotoxins in whole-animal systems is desirable in the context of enhancing the understanding of the pathogenesis of these toxins and exploring their uses as therapeutics. The development of such imaging assays is complicated by the often low numbers of toxin molecules targeting individual cells, making toxin detection challenging, as well as by the difficulty of providing sufficient spatial resolution to discriminate *bona-fide* intoxication from non-productive association of the toxin with the target cell, such as cell surface retention or endosomal/lysosomal sequestration.

Here, we describe a novel assay that provides single-cell-resolution imaging of anthrax toxin intoxication in animals. The assay takes advantage of the potential of genetic read-out systems to provide near-unlimited signal amplification, in this case, entailing the activation of a fluorescent reporter gene by an anthrax toxin lethal factor-Cre fusion protein that is translocated to the cytoplasm through the anthrax protective antigen heptamer or octamer pore and subsequently imported to the nucleus. Intoxication in some organs could be detected as early as 12 h after systemic introduction of the toxin, and, as expected, the efficiency of intoxication varied with toxin dose and mode of administration. The plasma membrane illumination provided by the membrane-targeted eGFP and tdTomato proteins, when combined with the staining of nuclei obtained by the *in vivo* Hoechst labelling procedure, provided sufficient cellular and tissue morphology information to allow for the identification of intoxicated cells in unfixed and unprocessed organ slices. It should also be noted that although only excised organs were examined here, the assay should be adaptable for multi-photon intravital imaging of cellular intoxication with essentially no modifications needed.

The development of our novel imaging assay afforded the first opportunity to directly visualize CMG-2- and TEM-8-mediated anthrax toxin intoxication and thereby enhance our understanding of the the two receptors in the intoxication process. Previous genetic studies in mice have demonstrated that CMG-2 deficiency conferred far greater resistance to anthrax lethal toxin and *B. anthracis* spore exposure than TEM-8 deficiency and identified the heart and liver as key CMG-2-dependent targets for, respectively, anthrax lethal toxin and anthrax edema toxin (11,56). Assuming that both receptors were close to ubiquitously expressed in tissues, this was tentatively suggested to be a consequence of a more than 10-fold lower affinity of PA for TEM-8 than for CMG-2. These findings are compatible with the current imaging study, showing that TEM-8 was essentially unable to support the intoxication in the heart, liver, and lungs, despite repeated toxin exposure through multiple injections. It should be noted, however, that full intoxication in other organs, including the spleen and kidney, was dependent upon TEM-8 but not CMG-2. This uequivocally demonstrates that TEM-8 is a functional receptor for anthrax toxin *in vivo* despite its reported lower affinity for PA.

A curious discrepancy between the aforementioned genetic studies and our current imaging study pertains to the intestine: Genetic studies have definitively established intestinal epithelial cells as direct targets for anthrax edema toxin (11). Nevertheless, we were consistently unable to observe the intoxication in intestinal epithelium. Intestinal epithelial cells of the *mTmG*^+/0^ reporter mice have previously been shown to undergo recombination *in vivo* and express eGFP in response to constitutive or inducible Cre expression, showing that the *mTmG* transgene is not inherently refractory to Cre-mediated recombination in intestinal epithelium (55). An attractive explanation for this discrepancy pertains to the lack of toxicity of LFn-NLS-Cre/PA as compared to edema toxin. We speculate that edema toxin may initially be unable to access this barrier tissue, but because damage to other visceral organs progresses as a consequence of EF intoxication, endothelial and/or epithelial barrier breakdown may allow entrance to the intestinal epithelial cells.

The imaging assay described here is simple and robust, and, importantly, it does not require the handling of toxic proteins. Therefore, it should be amenable for and adaptable to diverse research settings. The assay should be useful for answering a number of basic research questions regarding the pathogenicity of anthrax toxins, as well as assisting clinical/translational efforts aimed at optimizing the treatment of individuals accidentally or deliberately exposed to *Bacillus anthracis*.

Considerable effort is currently being expended on the development of modified anthrax toxins as novel agents for the treatment of human malignancies. Strategies employed include the reengineering of PA to bind tumor cell surface-enriched proteins (34,35) and the reengineering of PA to be proteolytically activated by proteases enriched in the tumor microenvironment, including matrix metalloproteinases (36–43), urokinase plasminogen activator (38,40,42,44–49), and testisin (50). The assay described here is imminently suited for assessing the efficiency of LF delivery to tumor-relevant cell populations by these modified PAs, as well as to systematically delineate off-targets, which may be invaluable for dose and route of delivery optimization. Last, but not least, by using PA variants selectively cleaved by specific cell surface proteases (43,46,50), the assay may be used for *in vivo* imaging of specific cell surface proteolytic activity in diverse physiological and pathological settings.

## ACKNOWLEDGEMENTS

We thank Annika Schmidt and Pia Mittweg for technical assistance. We thank Qian Ma for assistance with mouse work. We thank Dr. Mary Jo Danton for critically reviewing this manuscript. Supported by the NIAID (S.H.L) and NIDCR Intramural Research Program (T.H.B) of the NIH and the NIDCR Combined Technical Research Core (ZIC DE000729-09) and NIDCR Imaging Core: ZIC DE000750-01. RN, SN, and DP were funded by the European Union’s Horizon 2020 research and innovation program grant agreement No. 686841 (MAGNEURON).

