## Supplemental figures for "Direct imaging of anthrax intoxication in animals reveals shared and individual functions of CMG-2 and TEM-8 in cellular toxin entry"

[1]Proteases and Tissue Remodeling Section, National Institute of Dental and Craniofacial Research, National Institutes of Health, Bethesda, MD 20892, USA. [2]Laboratory of Parasitic Diseases, National Institute of Allergy and Infectious Diseases, National Institutes of Health, Bethesda, Maryland 20892, USA. [3]Department of Biochemistry II – Molecular Neurobiochemistry, Faculty of Chemistry and Biochemistry, Ruhr-Universität Bochum, 44801 Bochum, Germany.

[#]The two authors contributed equally to this research.

*Running title: *Visualizing anthrax toxin intoxication in animals*

Address correspondence and reprint requests to: Thomas H. Bugge, Ph.D., Proteases and Tissue Remodeling Section, National Institute of Dental and Craniofacial Research, National Institutes of Health, 30 Convent Drive, Room 211, Bethesda, MD 20892, Phone: (301) 827-4804, Fax: (301) 402-0823,

**Keywords:** Anthrax toxin/Cre recombinase/Cre reporter mice/CMG-2/confocal microscopy/TEM-8/ whole-animal imaging

---

### SUPPLEMENTAL FIGURE LEGENDS

**Supplemental Figure 1**. Amino acid sequences of the four LFn-Cre fusion proteins used in this study.

**Supplemental Figure 2.** Imaging mTomato fluorescence in slices of freshly excised organs from *mTmG*$^{+/0}$ transgenic mice by confocal microscopy. Low-magnification confocal images of fresh slices of the brain (A, A'), lungs (B, B'), testes (C, C'), large intestine (D, D'), lymph nodes (E, E'), thymus (F, F'), heart (G, G'), skin (H, H'), liver (I, I'), kidney (J, J'), spleen (K, K'), and trachea (L, L') from wildtype mice (A-L) and *mTmG*$^{+/0}$ mice (A'-L'). The images were assembled from individual images collected using a 20x objective and were stitched to cover the entire organ slice. Size bars: 1 mm.

**Supplemental Figure 3.** Imaging anthrax toxin intoxication in animals by using PA/LFn-NLS-Cre and *mTmG*$^{+/0}$ mice. Low-magnification confocal images of fresh slices of the heart (A-C'), kidney (D-F'), liver (G-I'), lungs (J-L'), and spleen (M-O') from *mTmG*$^{+/0}$ mice injected intravenously with 25 µg LFn-NLS-Cre alone (A, A', D, D', G, G', J, J', M, M'), 25 µg PA alone (B, B', E, E', H, H', K, K', N, N'), or with 25 µg LFn-NLS-Cre and 25 µg PA (C, C', F, F', I, I', L, L', O, O'). The images were assembled from individual images collected using a 20x objective and were stitched to cover the entire organ slice. Size bars: 1 mm.

**Supplemental Figure 4**. Absence of green fluorescence in organs from wildtype mice injected with 25 μg LFn-NLS-Cre and 25 μg PA. Low-magnification confocal images of fresh slices of the heart (A, B), kidney (C, D), liver (E, F), lungs (G, H), and spleen (I, J) from *mTmG*$^{+/0}$ mice (A, C, E, G, I) and wildtype mice (B, D, F, H, J) injected intravenously with 25 μg LFn-NLS-Cre and 25 μg PA. The images were assembled from individual images collected using a 20x objective and were stitched to cover the entire organ slice. Size bars: 1 mm. Note that panels A, C, E, G, and H are identical to, respectively, panels C', F', I', L', and O' in Supplemental Figure 3. Data presented in Supplemental Figure 3 and Supplemental Figure 4 are derived from one experiment.

**Supplemental Figure 5.** eGFP fluorescence is weak or absent in the large intestine, skin, and brain from mice injected with 25 µg LFn-NLS-Cre and 25 µg PA. Low-magnification confocal images of fresh slices of the large intestine (A, B), skin (C, D), and brain (E, F) from $mTmG^{+/0}$ mice injected intravenously with 25 µg LFn-NLS-Cre and 25 µg PA. The images were assembled from individual images collected using a 20x objective and were stitched to cover the entire organ slice. Size bars: 1 mm.

**Supplemental Figure 6.** Intravenously injected toxin disseminates more efficiently than intraperitoneally injected toxin. Low-magnification confocal images of fresh slices of the heart (A-D), kidney (E-H), liver (I-L), lungs (M-P), and spleen (Q-T) of $mTmG^{+/0}$ mice injected intraperitoneally (IP) with 25 μg LFn-NLS-Cre and 25 μg PA (A, E, I, M, Q), 50 μg LFn-NLS-Cre and 50 μg PA (B, F, J, N, R), or 75 μg LFn-NLS-Cre and 75 μg PA (C, G, K, O, S) or $mTmG^{+/0}$ mice injected intravenously (IV) with 25 μg LFn-NLS-Cre and 25 μg PA (D, H, L, P, T). The images were assembled from individual images collected using a 20x objective and were stitched to cover the entire organ slice. Size bars: 1 mm.

**Supplemental Figure 7.** eGFP expression increases with the number of toxin injections as well as with time after LFn-NLS-Cre and PA injection. Low-magnification confocal images of fresh slices of the heart (A-D'''), kidney (E-H'''), liver (I-L'''), lungs (M-P'''), and spleen (Q-T''') from *mTmG*$^{+/0}$ mice injected intravenously with 25 µg LFn-NLS-Cre and PA only on day zero (A-B', C, C', D, D', E-F', G, G', H, H', I-J', K, K', L, L', M-N', O, O', P, P', Q-R', S, S', T, T', single) or every day for 1, 2, 3, 4, or 5 days (B'', B''', C'', C''', D'', D''', F'', F''', G'', G''', H'', H''', J'', J''', K'', K''', L'', L''', N'', N''', O'', O'''. P'', P''', R'', R''', S'', S''', T'', T''', daily). Mice were imaged 24 h (A, A', E, E', I, I', M, M', Q, Q'), 48 h (B-B''', F-F''', J-J''', N-N''', R-R'''), 72 h (C-C''', G-G''', K-K''', O-O''', S-S'''), and 120 h (D-D''', H-H''', L-L''', P-P''', T-T''') after initial injection. The images were assembled from individual images collected using a 20x objective and were stitched to cover the entire organ slice. Size bars: 1 mm.

**Supplemental Figure 8.** eGFP expression in response to LFn-NLS-Cre and PA administration is dose-dependent. Low-magnification confocal images of fresh slices of the heart (A-C'), kidney (D-F'), liver (G-I'), lungs (J-L'), and spleen (M-O') from *mTmG*$^{+/0}$ mice injected intravenously with 5 µg LFn-NLS-Cre and 5 µg PA (A, A', D, D', G, G', J, J', M, M'), 15 µg LFn-NLS-Cre and 15 µg PA (B, B', E, E', H, H', K, K', N, N'), or 25 LFn-NLS-Cre and 25 µg PA (C, C', F, F', I, I', L, L', O, O'). The images were assembled from individual images collected using a 20x objective and were stitched to cover the entire organ slice. Size bars: 1 mm.

**Supplemental Figure 9.** eGFP expression can be detected early after LFn-NLS-Cre and PA administration. Low-magnification confocal images of fresh slices of the heart (A-D'), kidney (E-H'), liver (I-L'), lungs (M-P'), and spleen (Q-T') from *mTmG*$^{+/0}$ mice analyzed 6 h (A, A', E, E', I, I', M, M', Q, Q'), 8 h (B, B', F, F', J, J', N, N', R, R'), 10 h (C, C',G, G', K, K', O, O', S, S'), and 12 h (D, D', H, H', L, L', P, P', T, T') after intravenous administration of 75 µg LFn-NLS-Cre and 75 µg PA. The images were assembled from individual images collected using a 20x objective and were stitched to cover the entire organ slice. Size bars: 1 mm.

**Supplemental Figure 1.**

Sequences of the four LFn-Cre fusion proteins used in this study.

**LFn-NLS-Cre:** SUMO domain is in red, the LFn in blue, the NLS in green, and Cre in black

MGSSHHHHHHGSGLVPRGSASMSDSEVNQEAKPEVKPEVKPETHINLKVSDGSSEIFFKIKKTTPLRRLMEAFAKRQGKEMDSLRFLYDGIRIQADQTPEDLDMEDNDII
EAHREQIGGAGGHGDVGMHVKEKEKNKDENKRKDEERNKTQEEHLKEIMKHIVKIEVKGEEAVKKEAAEKLLEKVPSDVLEMYKAIGGKIYIVDGDITKHISLEALSED
KKKIKDIYGKDALLHEHYVYAKEGYEPVLVIQSSEDYVENTEKALNVYYEIGKILSRDILSKINQPYQKFLDVLNTIKNASDSDGQDLLFTNQLKEHPTDFSVEFLEQNSN
EVQEVFAKAFAYYIEPQHRDVLQLYAPEAFNYMDKFNEQEINLGGGSKKKRKVSNLLTVHQNLPALPVDATSDEVRKNLMDMFRDRQAFSEHTWKMLLSVCRSWAA
WCKLNNRKWFPAEPEDVRDYLLYLQARGLAVKTIQQHLGQLNMLHRRSGLPRPSDSNAVSLVMRRIRKENVDAGERAKQALAFERTDFDQVRSLMENSDRCQDIRNL
AFLGIAYNTLLRIAEIARIRVKDISRTDGGRMLIHIGRTKTLVSTAGVEKALSLGVTKLVERWISVSGVADDPNNYLFCRVRKNGVAAPSATSQLSTRALEGIFEATHRLIY
GAKDDSGQRYLAWSGHSARVGAARDMARAGVSIPEIMQAGGWTNVNIVMNYIRNLDSETGAMVRLLEDGD

The 3 other proteins that were made and compared differed only in the linker region.
An alignment of the four proteins in this region is shown here:
NLS is in green and bold
Ubiquitin is underlined in LFn-modified ubiquitin-NLS-Cre
The ubiquitin mutations in LFn-modified ubiquitin-NLS-Cre are in blue, and include V5A, V17A, and all 7 K to R

| | |
|---|---|
| **LFn-NLS-Cre:** | NYMDKFNEQEINLGGG------------------------------------------ |
| **LFn-Cre:** | NYMDKFNEQEINLGGGSS---------------------------------------- |
| **LFn-native ubiquitin-NLS-Cre:** | NYMDKFNEQEINLGGGSMQIFVKTLTGKTITLEVEPSDTIENVKAKIQDKEGIPPDQQRL |
| **LFn-modified ubiquitin-NLS-Cre:** | NYMDKFNEQEINLGGGSMQIF**AR**TLTG**R**TITLE**A**EPSDTIENV**R**A**R**IQD**R**EGIPPDQQRL |

| | |
|---|---|
| **LFn-NLS-Cre:** | ---------------------------------S**KKKRKV**SNLLTVHQNLPALPVDATS |
| **LFn-Cre:** | ----------------------------------------NLLTVHQNLPALPVDATS |
| **LFn-native ubiquitin-NLS-Cre:** | IFAGKQLEDGRTLSDYNIQKESTLHLVLRLRGGGS**KKKRKV**SNLLTVHQNLPALPVDATS |
| **LFn-modified ubiquitin-NLS-Cre:** | IFAG**R**QLEDGRTLSDYNIQ**R**ESTLHLVLRLRGGGS**KKKRKV**SNLLTVHQNLPALPVDATS |

Supplemental Figure 1

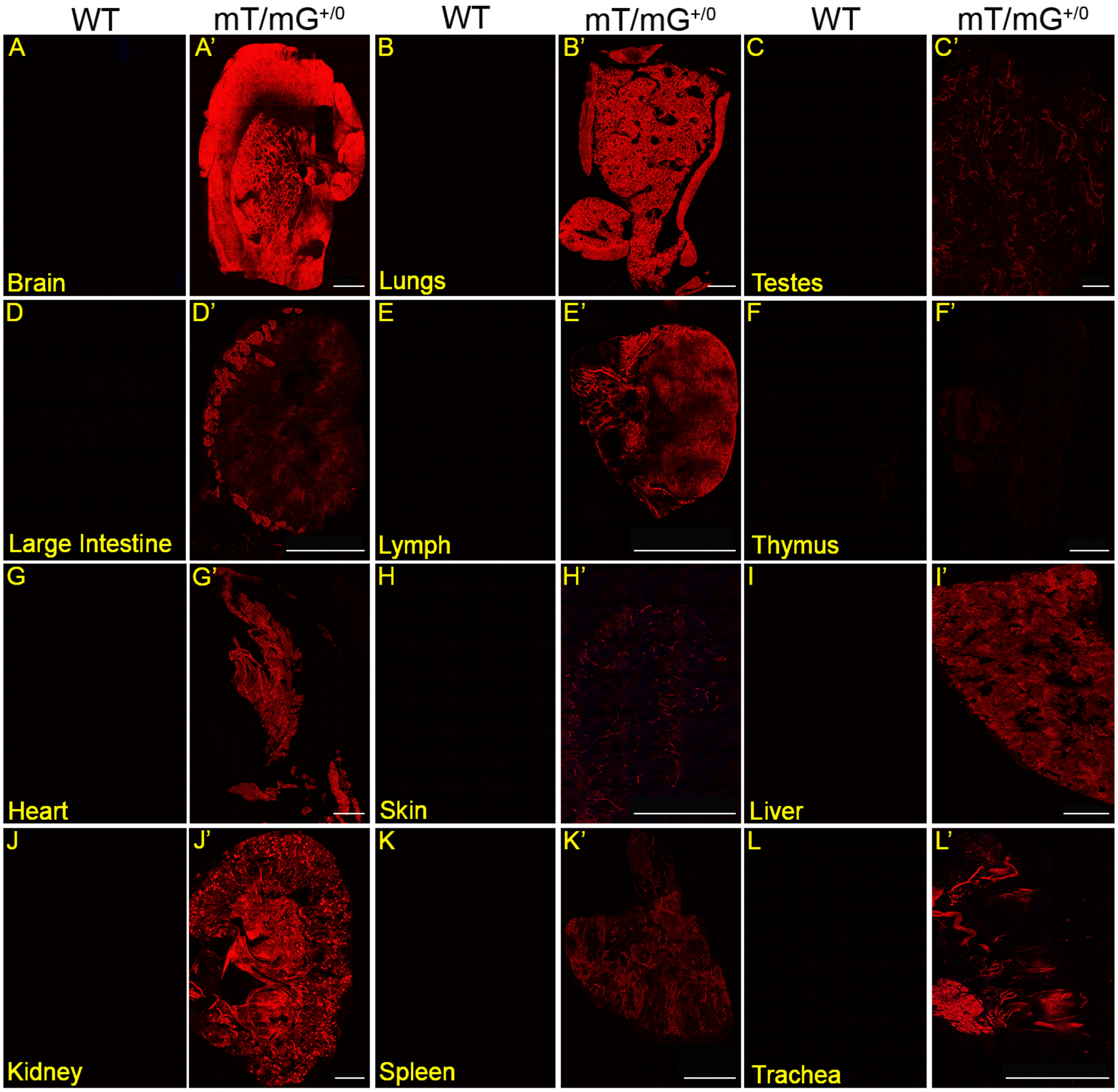


Supplemental Figure 2

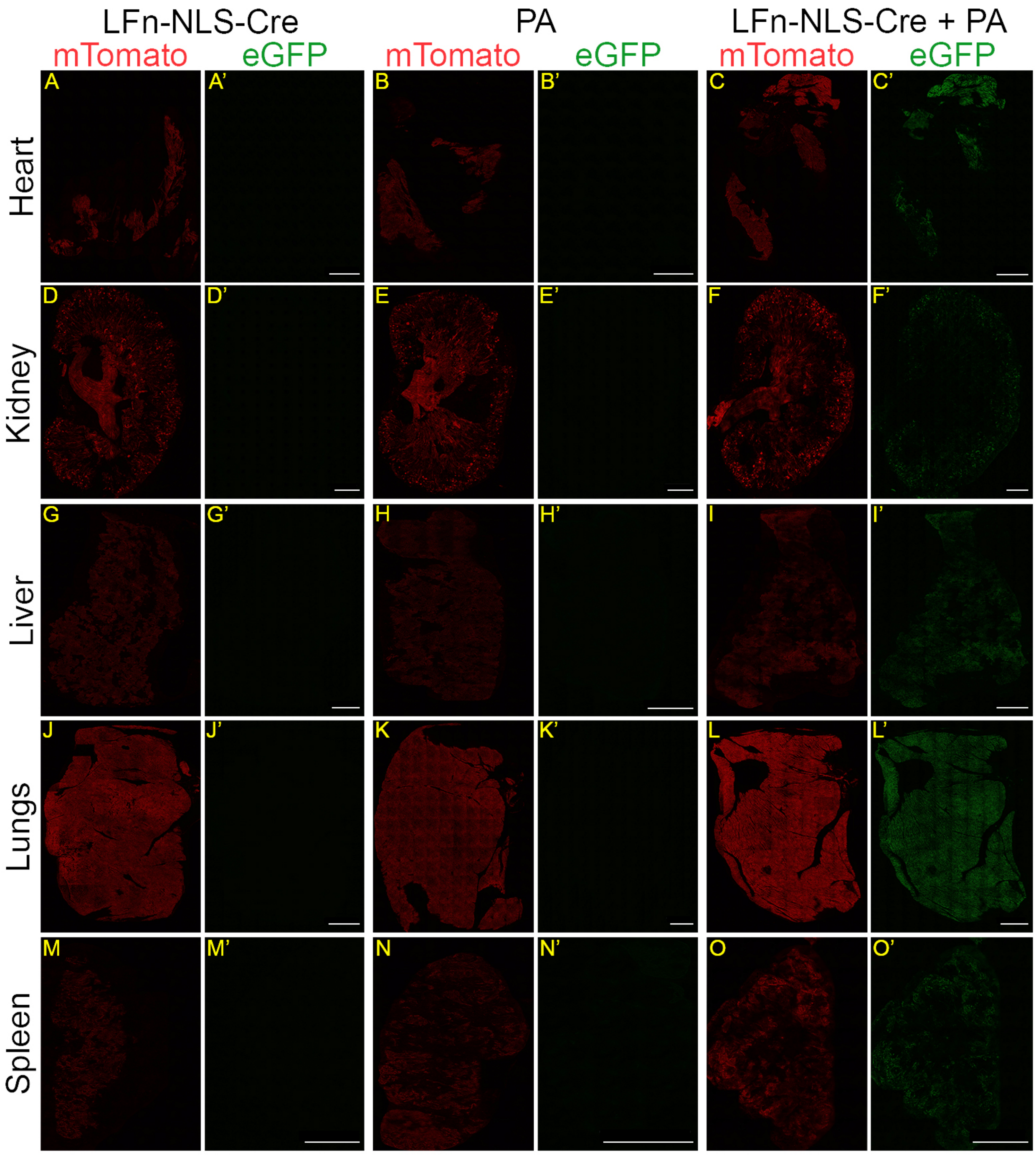


Supplemental Figure 3

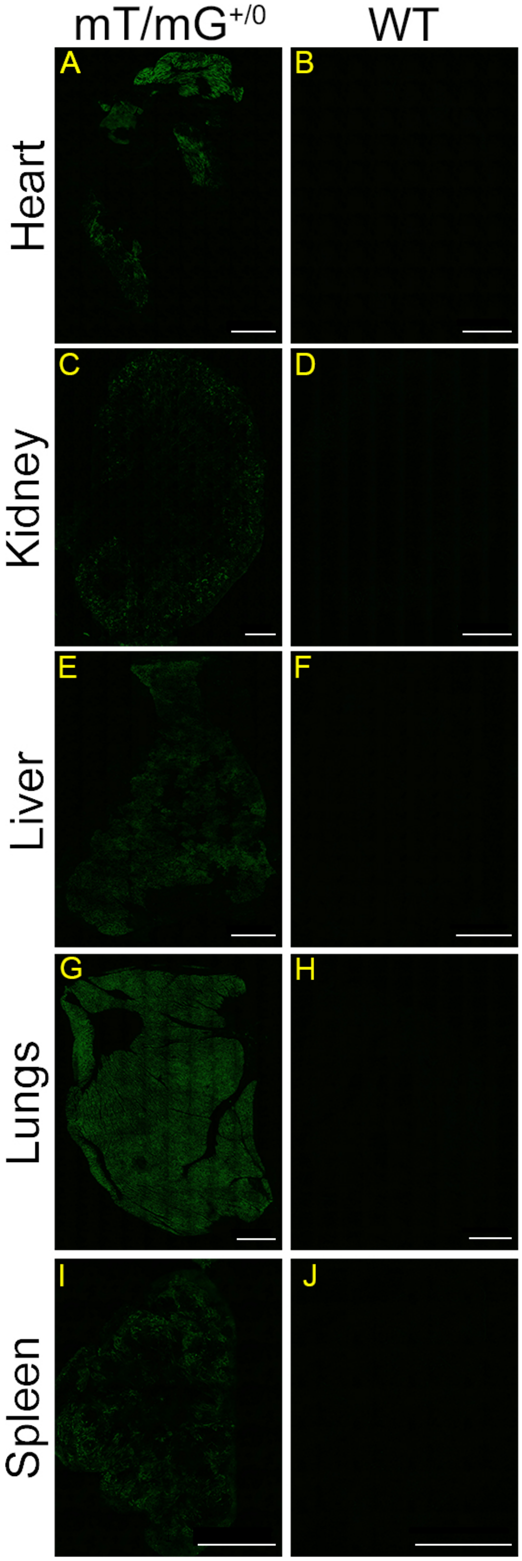


Supplemental Figure 4

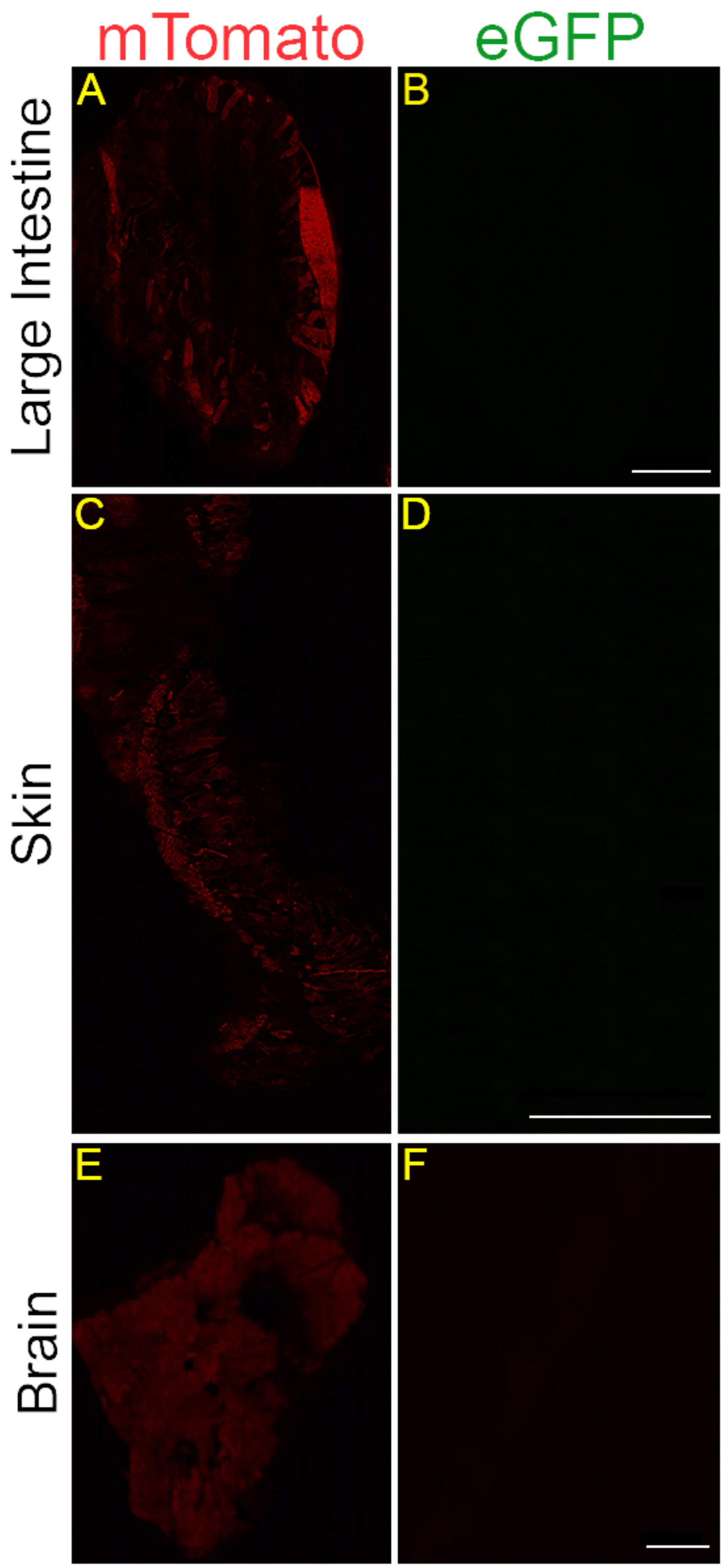


Supplemental Figure 5

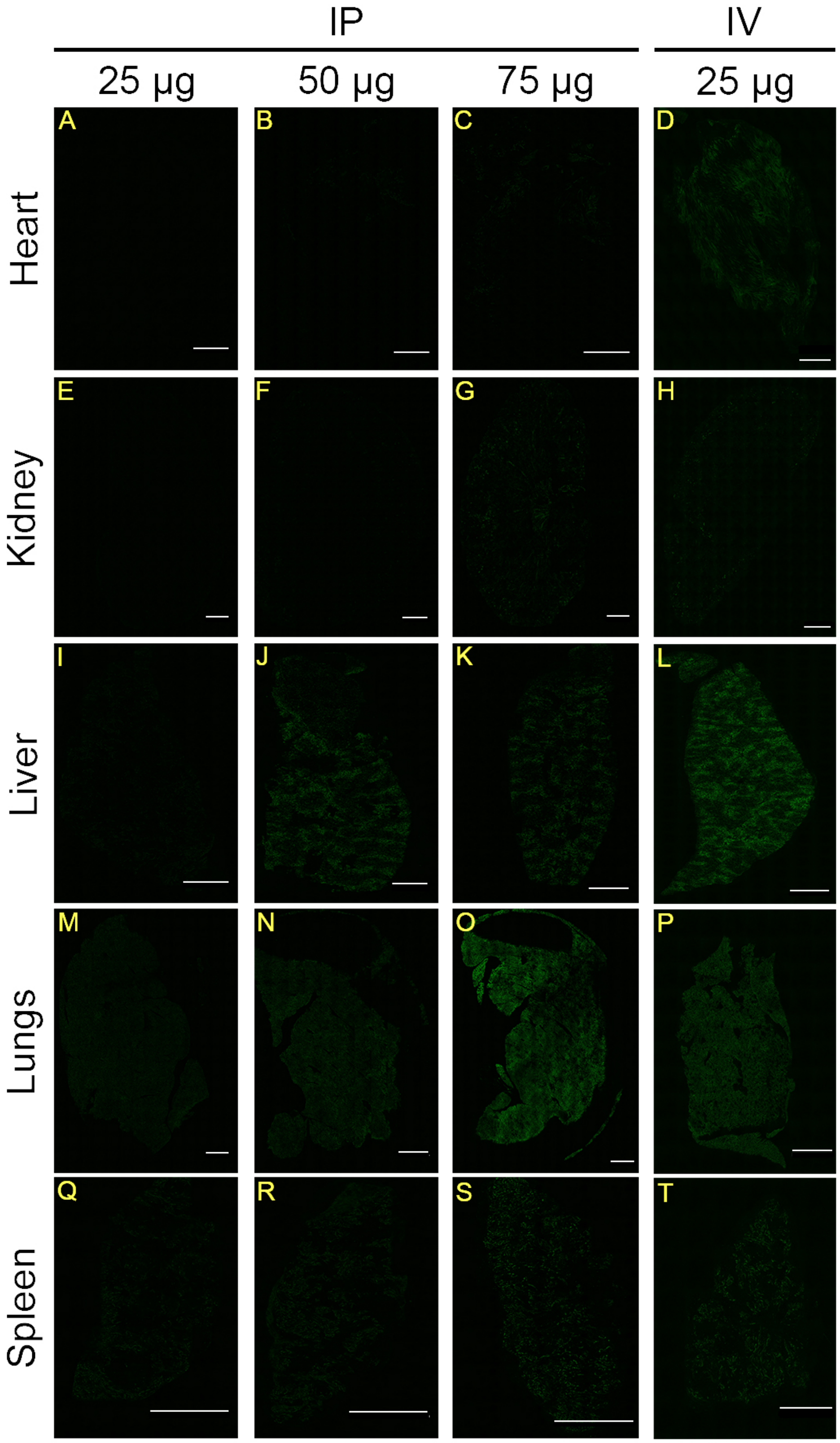


Supplementary Figure 6

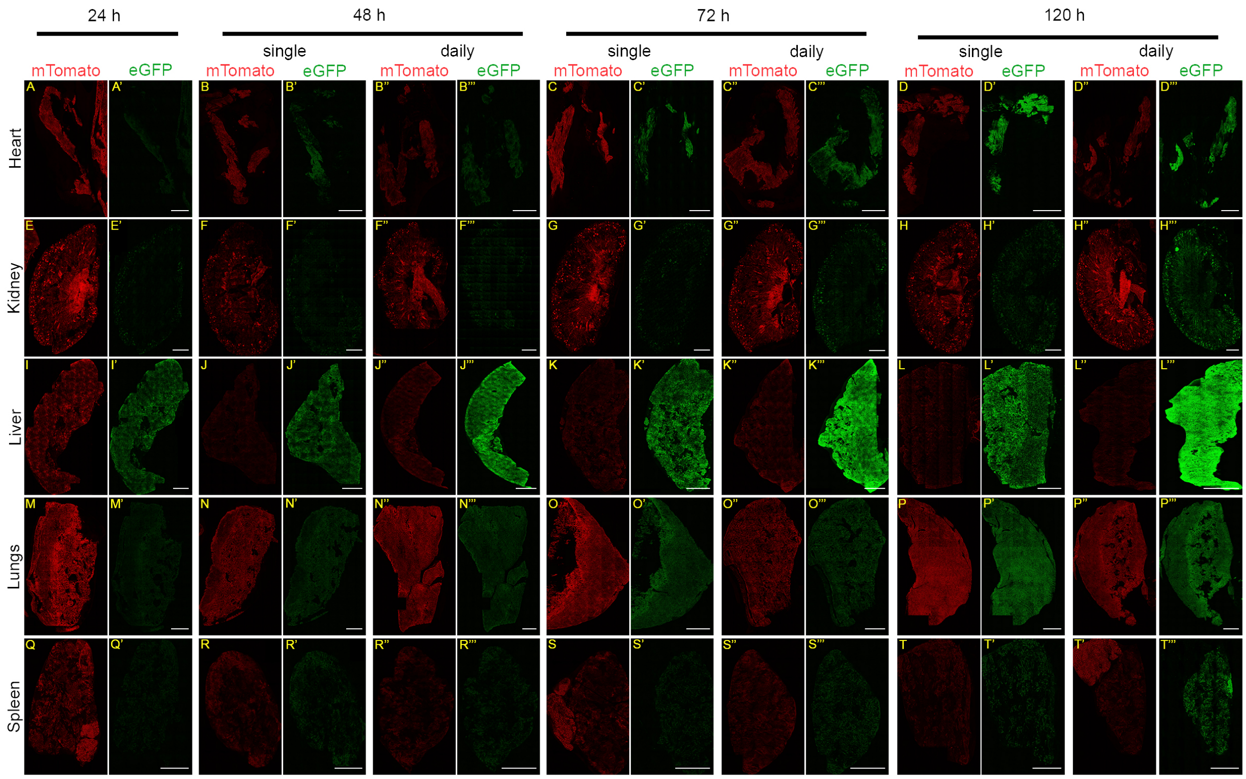


Supplementary Figure 7

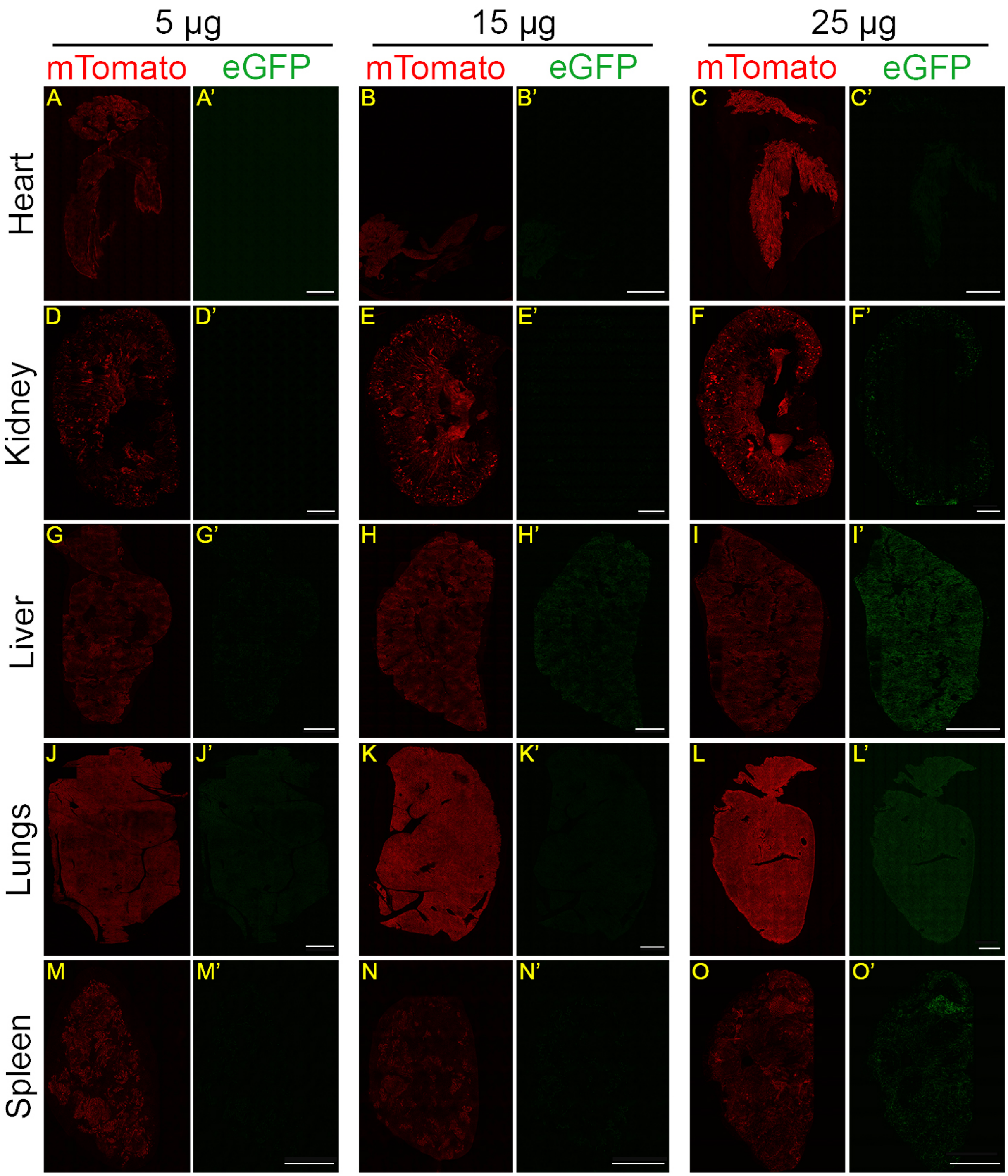


Supplementary Figure 8

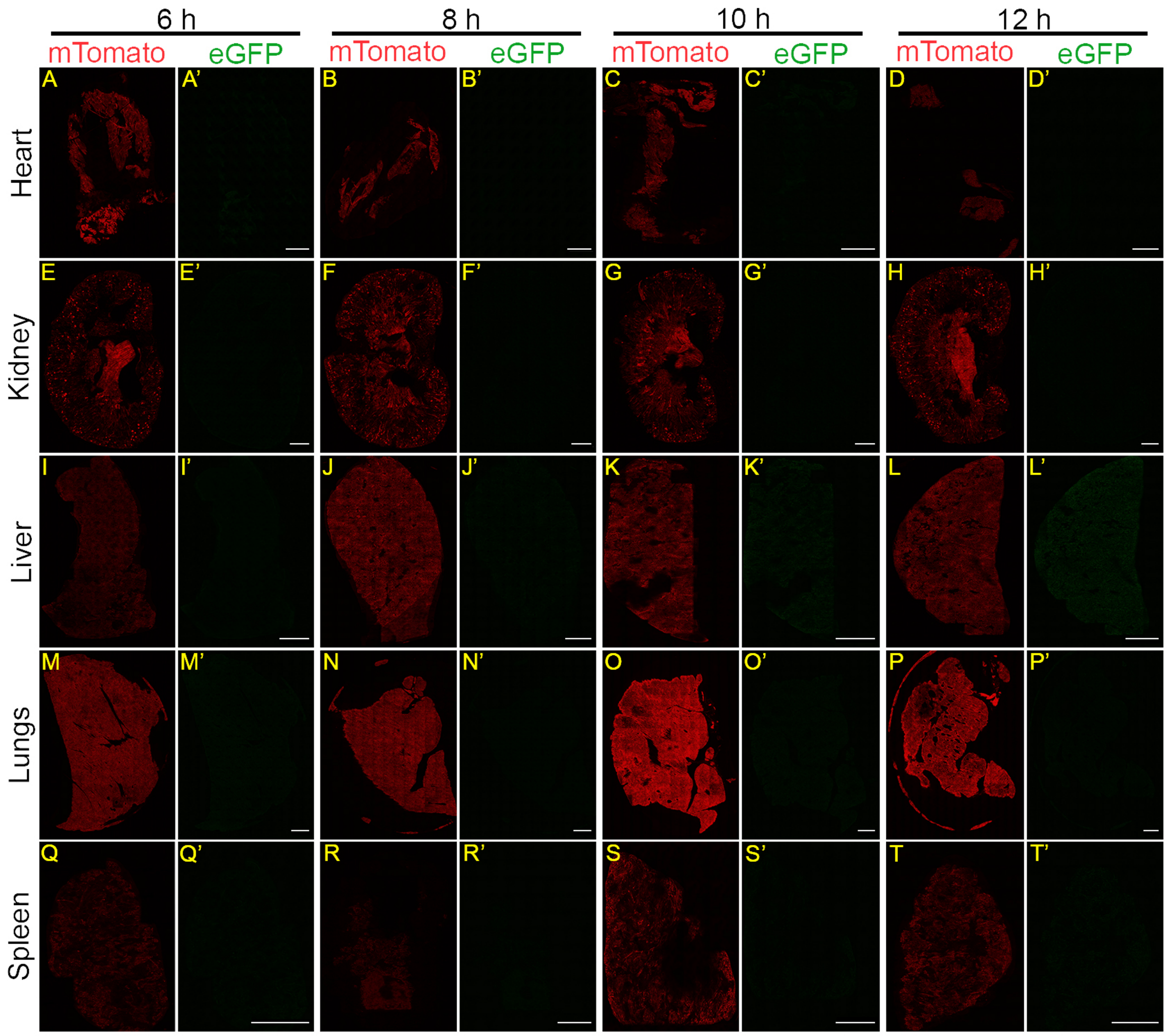


Supplementary Figure 9
